# Photo-inactivation of *Neisseria gonorrhoeae*: A paradigm changing approach for combating antibiotic-resistant gonococcal infection

**DOI:** 10.1101/422634

**Authors:** Ying Wang, Raquel Ferrer-Espada, Yan Baglo, Xueping S. Goh, Kathryn D. Held, Yonatan H. Grad, Ying Gu, Jeffrey A. Gelfand, Tianhong Dai

## Abstract

*Neisseria gonorrhoeae* is the causative pathogen of the sexually transmitted disease gonorrhea, a disease at risk of becoming untreatable due to increasing antibiotic resistance. There is a critical need for the development of new anti-gonococcal therapies. In this study, we investigated the effectiveness of antimicrobial blue light (aBL), an innovative non-antibiotic approach, for the inactivation of antibiotic-resistant *N*. *gonorrhoeae*. Our findings indicated that aBL at 405 nm preferentially inactivated antibiotic-resistant *N. gonorrhoeae* over the vaginal epithelial cells. Furthermore, no genotoxicity of aBL to the vaginal epithelial cells was observed at the exposure for inactivating *N. gonorrhoeae*. aBL also effectively inactivated *N. gonorrhoeae* that had invaded into the vaginal epithelial cells. No gonococcal resistance to aBL developed after 15 successive cycles of sub-therapeutic inactivation. Endogenous aBL-active photosensitizing chromophores (porphyrins and flavins) in *N. gonorrhoeae* were identified and quantified using ultra performance liquid chromatography, with coproporphyrin being the most abundant endogenous porphyrin species. Taken together, aBL at 405 nm represents a potent potential treatment for gonococcal infections.

**One Sentence Summary:** aBL selectively inactivated antibiotic-resistant *Neisseria gonorrhoeaen* over normal vaginal epithelial cells.

## INTRODUCTION

*Neisseria gonorrhoeae* is a human-specific bacterial pathogen and the cause of gonococcal infection (gonorrhea) (1, 2). If not appropriately treated, gonorrhea can result in severe sequelae such as pelvic inflammatory disease, infertility, ectopic pregnancy, first trimester abortion, neonatal conjunctivitis leading to blindness and, less frequently, male infertility and disseminated gonococcal infections (3). The CDC estimates that 820,000 new gonococcal infections occur each year in the United States (4), with the estimated total annual economic cost for the treatment of gonorrhea approaching $5 billion (5). Furthermore, gonorrhea facilitates the transmission and acquisition of HIV (6-8). In the absence of a gonococcal vaccine, public health control of gonorrhea relies primarily on the availability of effective antibiotic treatment (3). However, *N*. *gonorrhoeae* has become resistant to each of the first-line antibiotics that have been used to treat it (9, 10). The antibiotic combination of azithromycin and ceftriaxone is the current CDC recommendation for treating gonorrhea (11). Worryingly, the susceptibility to ceftriaxone has been decreasing globally in recent years, resistance to azithromycin is prevalent and emerges rapidly in settings where this drug is frequently used, and *N*. *gonorrhoeae* strains with decreased susceptibility or resistance to ceftriaxone and concomitant azithromycin resistance are already circulating globally (12). For example, a cluster of cases was recently reported in Hawaii that was resistant to both azithromycin and ceftriaxone (13, 14). The CDC flagged multidrug-resistant *N*. *gonorrhoeae* as an urgent threat to human health (15), the highest level of concern.

As the use of antibiotics drives the emergence and spread of multidrug-resistant *N*. *gonorrhoeae* (3, 16-18), there is a critical need for the development of new treatment strategies (19, 20). An innovative non-antibiotic therapy, antimicrobial blue light (aBL), has attracted increasing attention due to its intrinsic antimicrobial properties without the involvement of exogenous photosensitizers (21, 22). Although the mechanism of action of aBL is still not fully understood, a common hypothesis is that aBL excites naturally occurring endogenous photosensitizers (e.g., iron-free porphyrins or/and flavins) within bacterial cells and subsequently leads to the production of cytotoxic oxidative species (23, 24).

The advantages of aBL over traditional antibiotics include rapid action and equal inactivation effectiveness independent of antibiotic resistance (21, 22). It is also currently thought that bacteria are less able to develop resistance to aBL than to traditional antibiotics because of the multi-target characteristic of aBL (21, 22, 25). In comparison to traditional antimicrobial photodynamic therapy, aBL is appealing in that it inactivates bacteria (renders bacteria unable to multiply) without the involvement of exogenous photosensitizers (26-32). In addition, it is well accepted that aBL is much less detrimental to the host cells than germicidal ultraviolet irradiation (33, 34).

aBL has never been studied as a treatment for gonorrhea (35). In this study, we evaluated the effectiveness of aBL in rendering antibiotic-resistant *N*. *gonorrhoeae* unable to multiply (termed inactivation), the toxicity of aBL to the epithelial cells, and the potential development of gonococcal resistance to aBL. In addition, we determined and quantified the presence of aBL-active endogenous photosensitizing chromophores in *N. gonorrhoeae*.

## RESULTS

### Effectiveness of aBL inactivation of *N. gonorrhoeae* in planktonic suspensions

To achieve 3-log_10_ CFU inactivation of *N. gonorrhoeae* in suspensions, aBL exposures of 25 J/cm^2^ (for ATCC700825; ∽7 min illumination) to 59 J/cm^2^ (for Strain #199; ∽16 min illumination) were used (Fig. 1A). To inactivate all the bacterial CFU in suspensions (> 6-log_10_ CFU inactivation), aBL exposures of 45 J/cm^2^ (for ATCC700825; 12.5 min illumination) to 108 J/cm^2^ (for Strain #199; 30 min illumination) were applied (Fig. 1A).

**Fig. 1.**
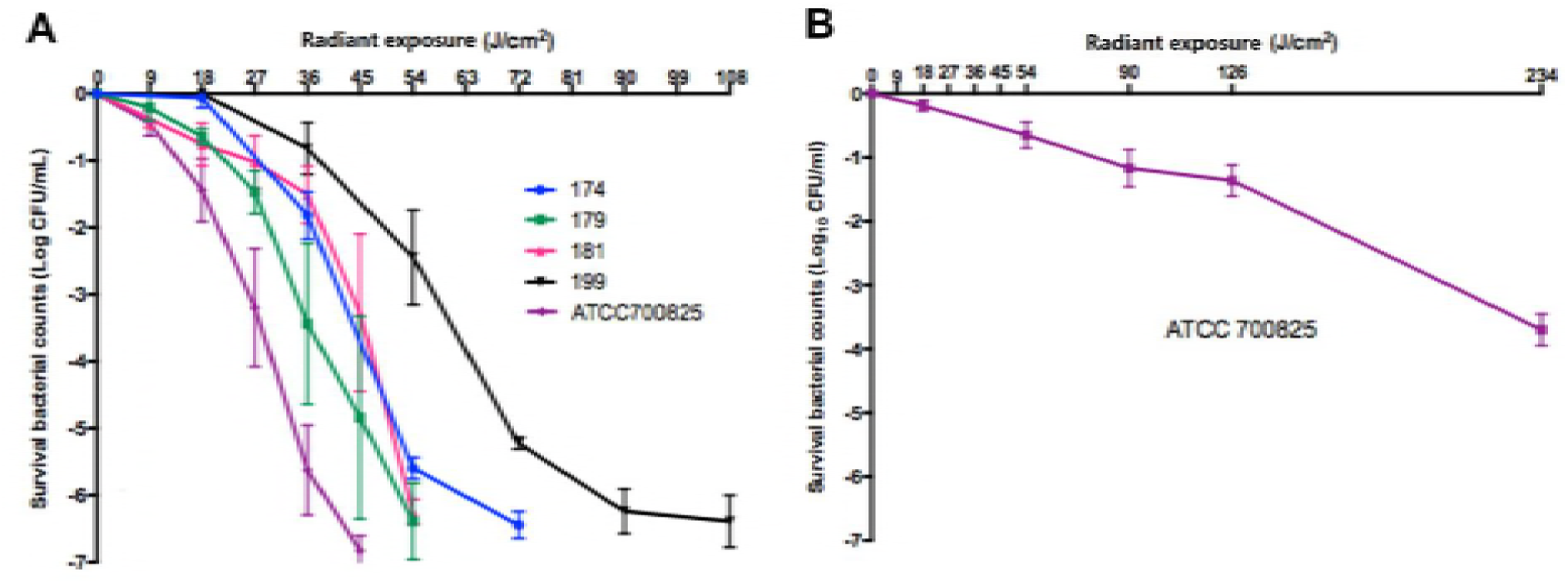
Effectiveness of aBL inactivation of *N. gonorrhoeae* at different wavelengths. (**A**) 405-nm wavelength. ATCC 700825 and clinical isolates 174, 179, 181, 199 were studied. **(B)** 470 nm wavelength. ATCC 700825 was studied. Bars: standard error of the means.

The anti-gonococcal effect of 470-nm aBL was tested on ATCC 700825, and was much less effective than 405-nm aBL (Fig. 1B). An aBL exposure of 170 J/cm^2^ (∽ 47 min illumination) was required to produce 3-log_10_ CFU inactivation. According to this finding, we employed 405-nm wavelength for the studies following.

Transmission electron microscopy (TEM) images taken after varying aBL exposures revealed aBL-induced morphological changes in *N. gonorrhoeae* cells, such as formation of bulges (black arrows), breakage of cell walls accompanied by cytoplasmic release (blue arrows), loss of outer membranes (white arrows) and cellular disintegration (blue stars) (Fig. 2). Decreased density of cytoplasm was observed in some cells accompanied by increased cellular size, suggesting that aBL induced damage to the cell wall of *N. gonorrhoeae*.

**Fig. 2.**
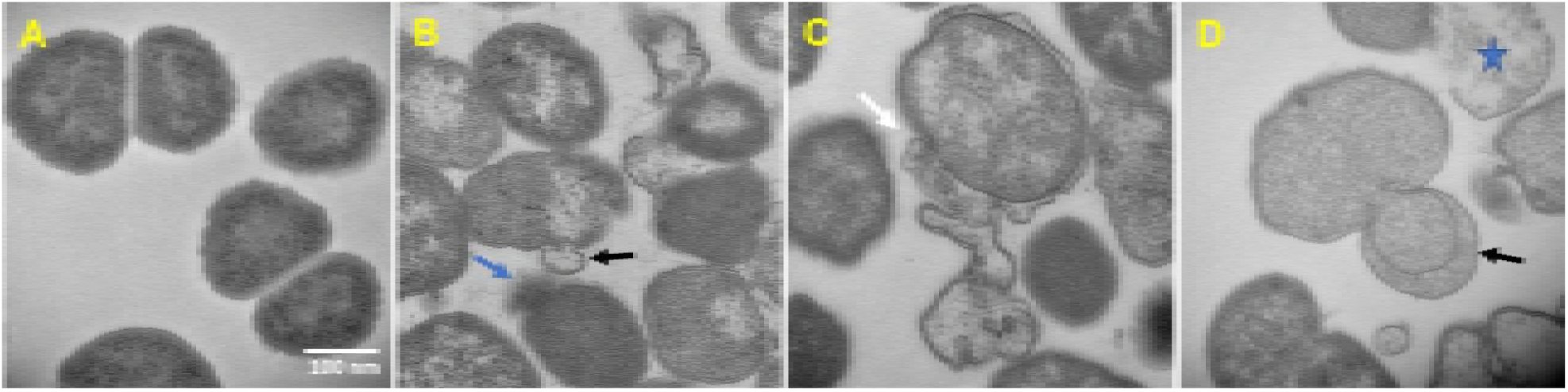
Transmission electron microscopy (TEM) images of untreated and 405-nm aBL treated *N. gonorrhoeae* (ATCC700825). **Panels A – D are the images of cultures exposed to 0 (no aBL), 9, 18, and 27 J/cm^2^ aBL, respectively. Images were taken immediately after each aBL exposure. Black arrows: bulges formation; blue arrows: breakage of cell wall accompanied by cytoplasmic release; white arrows: loss of outer membrane; blue stars: cellular disintegration. Scale bar: 100 nm**.

### Toxicity of 405-nm aBL to normal vaginal epithelial cells

As shown in Fig.1A, a 405-nm aBL exposure of < 108 J/cm^2^ resulted in complete eradication of *N. gonorrhoeae* in suspensions (> 6-log_10_ CFU inactivation). In contrast, the toxicity study of aBL using normal vaginal epithelial cells showed that no statistically significant loss of viability of VK2/E6E7 cells was observed when aBL exposures of up to 108 J/cm^2^ were applied (*P* =0.6676) (Fig. 3A). Comet assay results showed that no aBL induced DNA damage occurred in the VK2/E6E7 cells at aBL exposures up to 216 J/cm^2^ (Fig. 3B).

**Fig. 3.**
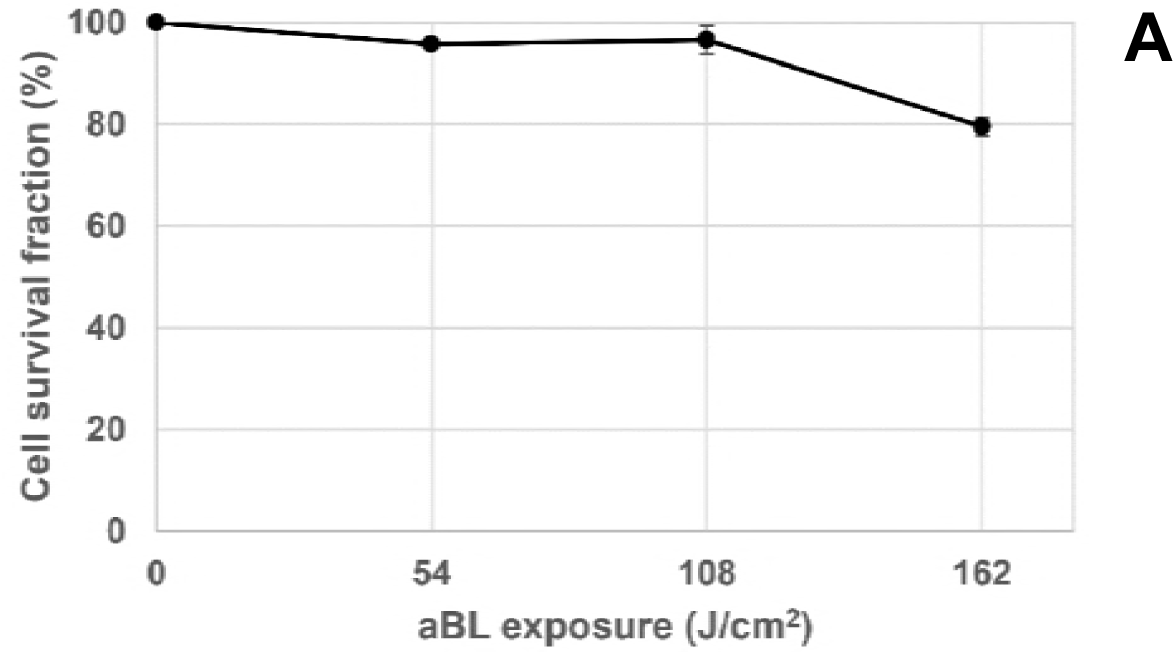
Toxicity of 405-nm aBL to normal vaginal epithelial cells. **(A)** Change of the viability of normal VK2/E6E7 cells in response to 405-nm aBL at different exposures. Viability of VK2/E6E7 cells: 0 J/cm^2^ vs. 54 J/cm^2^: *P* = 0.3474, 0 J/cm^2^ vs. 108: *P* = 0.6676; 0 J/cm^2^ vs. 162 J/cm^2^: *P*< 0.0001. Bars denote standard errors. (**B**) Comet assay images of normal VK2/E6E7 cells with and without 405-nm aBL exposure. The 0 J/cm^2^, 54 J/cm^2^, 108 J/cm^2^, and 216 J/cm^2^ images show the cultures exposed to 0 J/cm^2^, 9 J/cm^2^, 18 J/cm^2^, and 27 J/cm^2^ aBL, respectively. Positive control is treated with H_2_O_2_.

### Presence and quantity of endogenous porphyrins and flavins in *N. gonorrhoeae*

The UPLC chromatograms identified the presence of several types of endogenous porphyrins in *N. gonorrhoeae*, including uroporphyrin, 7-carboxylporphyrin, 6-carboxylporphyrin, 5-carboxylporphyrin and coproporphyrin (Fig. 4). No PpIX was detected in any of the five *N. gonorrhoeae* strains studied. Quantitative analysis revealed that the most abundant type of endogenous porphyrin species in *N. gonorrhoeae* was coproporphyrin, with concentrations ranging from 7.93 to 19.16 nmol/g (protein weight) in the five *N. gonorrhoeae* strains. The concentrations and content percentages of each porphyrin species in the *N. gonorrhoeae* strains are summarized in Table 1.

**Table 1.**
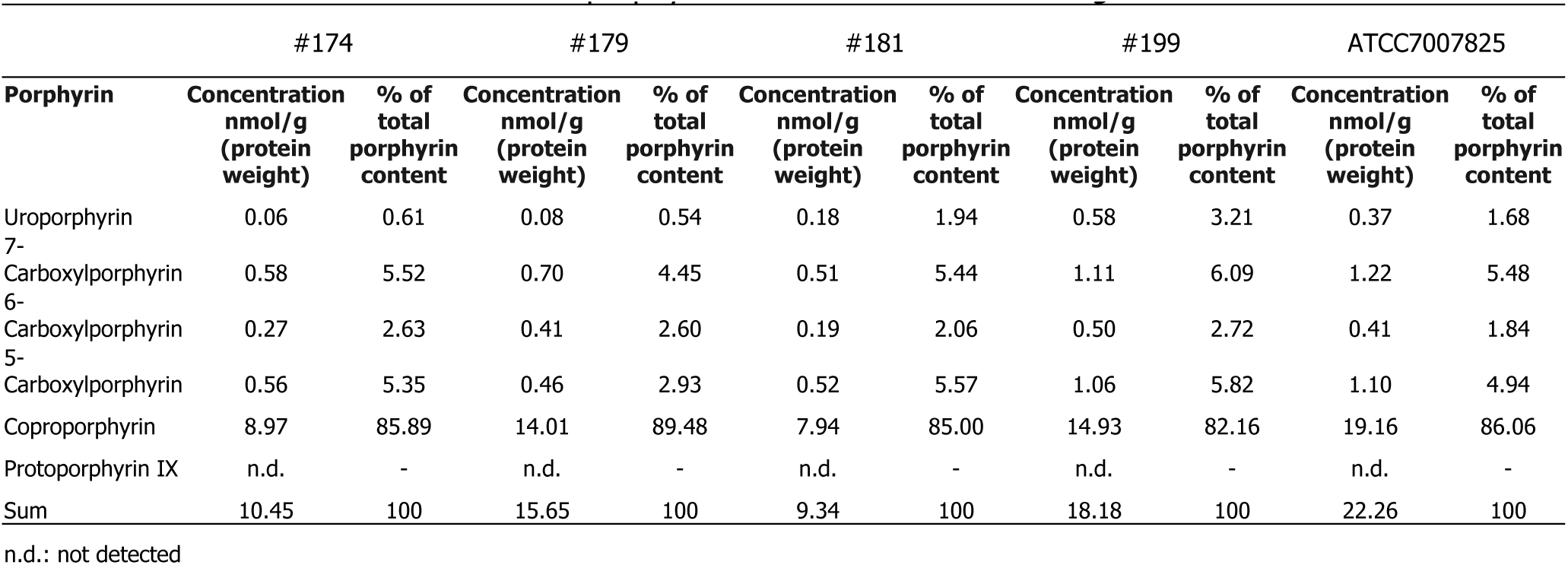
Concentrations of porphyrins in the extracts of five *N. gonorrhoeae* strains

**Fig. 4.**
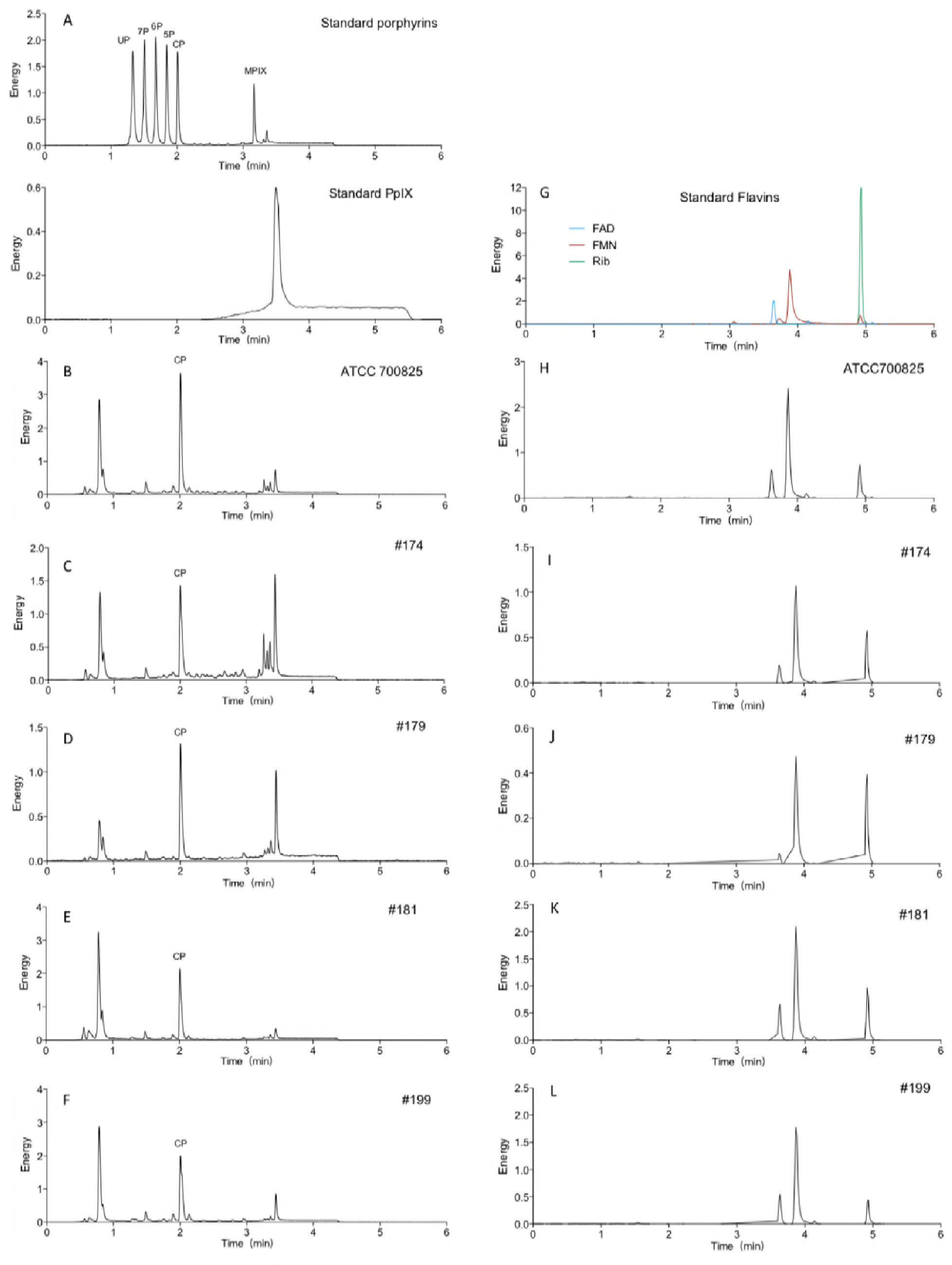
Chromatograms from the UPLC analysis. A: standard porphyrin and PPIX. UP: uroporphyrin I; 7P: 7-carboxylporphyrin I; 6P: 6-carboxylporphyrin I; 5P: 5-carboxylporphyrin I; CP: coproporphyrin; MPIX: mesoporphyrin IX; PPIX: protoporphyrin IX. B, C, D, E, and F: extracts of ATCC700825, #174, #179, #181, #199, respectively. G: standard flavins. H, I, J, K, and L: G: extracts of ATCC700825, #174, #179, #181, #199, respectively.

UPLC results also indicated the presence of flavins in *N. gonorrhoeae* (Fig. 5). For all the strains except #179, the concentrations of FAD and FMN were higher than that riboflavin (Fig. 5).

**Fig. 5.**
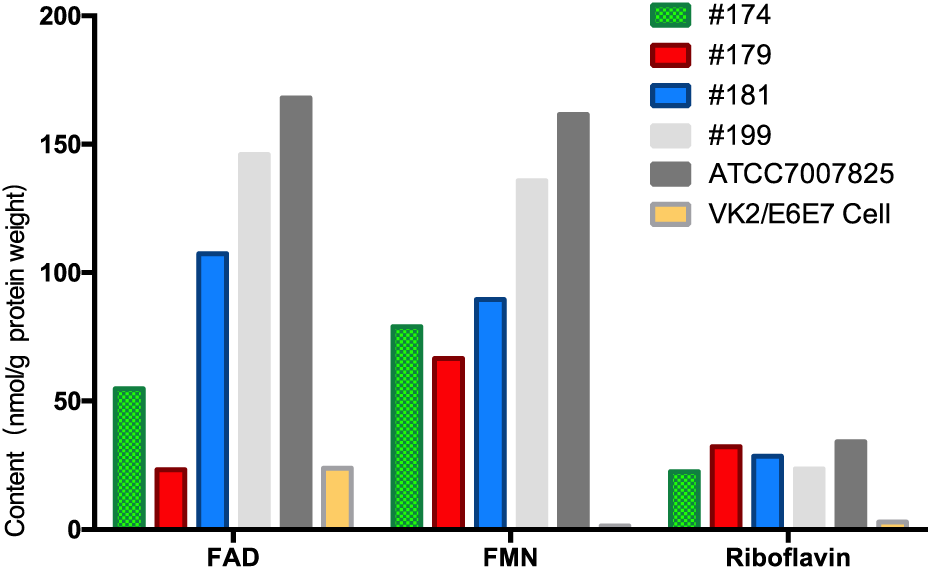
Concentrations of FAD, FMN, and Riboflavin in the extracts of *N. gonorrhoeae and* VK2/E6E7 cells.

### Presence and quantity of endogenous porphyrins and flavins in normal vaginal epithelial cells

Quantitative analysis by UPLC showed that the total content of endogenous porphyrins in VK2/E6E7 cells is 0.0176 nmol/g (protein weight) (Table 2), which is hundreds of times lower than the total content of porphyrins in *N. gonorrhoeae* (9.34 to 22.26 nmol/g; Table 1). The content of flavins in VK2/E6E7 cells is also significantly lower than that in the *N. gonorrhoeae* (Fig. 5).

**Table 2.** <u>**Concentrations of endogenous porphyrins in VK2/E6E7 cells**</u>

| Porphyrin | Concentration nmol/g (protein weight) | % of total porphyrin content |
| --- | --- | --- |
| Uroporphyrin | 0.0074 | 41.95 |
| 7-Carboxylporphyrin | n.d. | - |
| 6-Carboxylporphyrin | n.d. | - |
| 5-Carboxylporphyrin | n.d. | - |
| Coproporphyrin | 0.0102 | 58.05 |
| Protoporphyrin IX | n.d. | - |
| Sum | 0.0176 | 100 |

n.d.: not detected

### Potential development of gonococcal resistance to aBL

Fig. 8 shows the log_10_-CFU reduction of ATCC 700825 in response to 15 successive cycles of sub-therapeutic aBL exposures at 405 nm. Correlation analysis found a weak tendency of increased aBL inactivation of *N. gonorrhoeae* with the number of cycles (correlation coefficient =0.24). However, the correlation is not statistically significant (*P* =0.40), indicating that development of gonococcal resistance to aBL did not occur after 15 successive cycles of sub-therapeutic aBL exposures.

### Effect of bacterium-host cell interaction on the tolerance of the vaginal epithelial cells to aBL and the efficacy of aBL inactivation of *N. gonorrhoeae*

Fig. 7 displays the viability changes of VK2/E6E7 cells and *N. gonorrhoeae* cells in the co-cultures of VK2/E6E7 cells and *N. gonorrhoeae* in response to 405-nm aBL. In comparison with the result of non-infected VK2/E6E7 cells shown in Fig. 3, it can be seen that infected VK2/E6E7 cells are more susceptible to aBL inactivation than non-infected VK2/E6E7 cells.

**Fig. 6.**
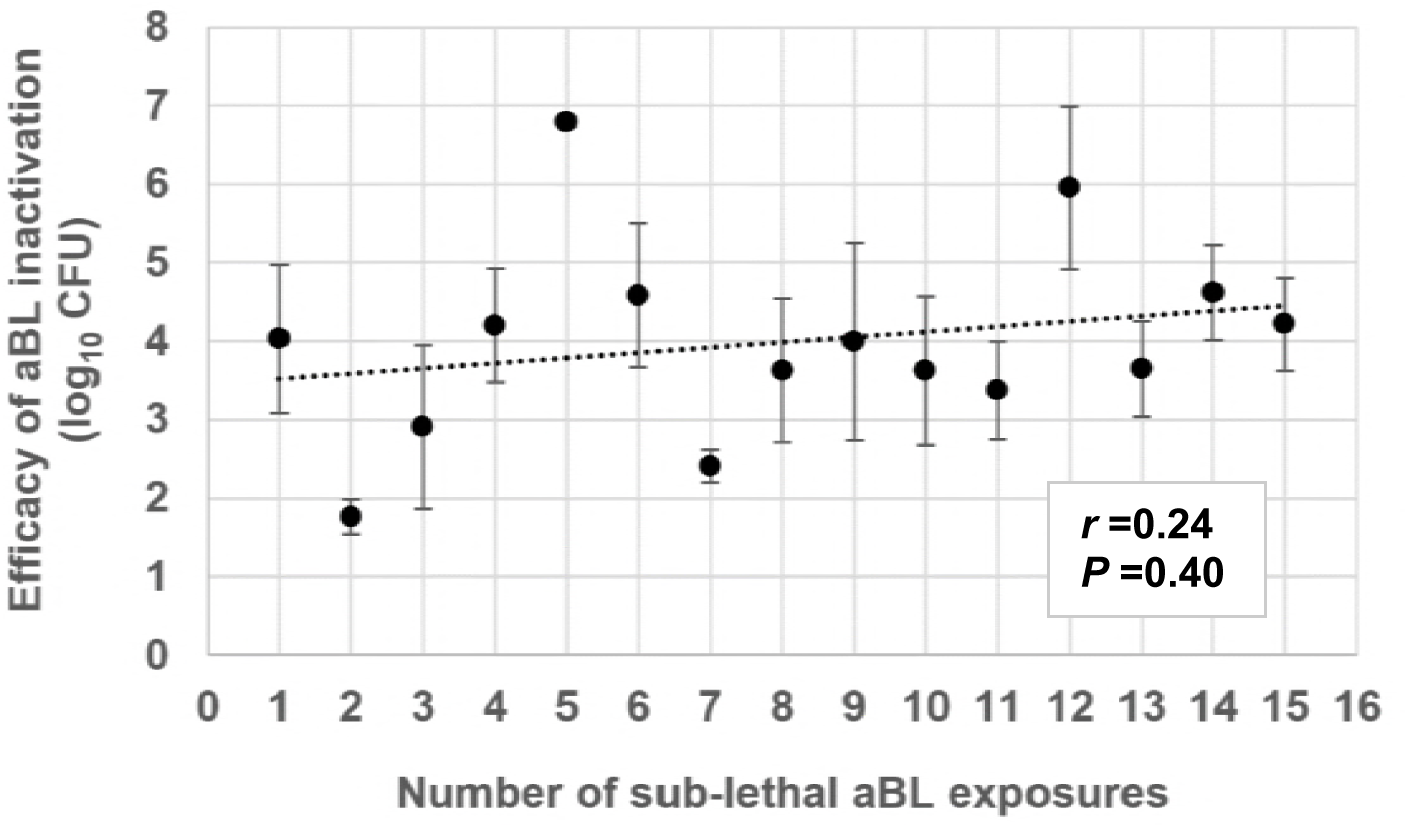
Efficacy of 405-nm aBL inactivation of *N. gonorrhoeae* (ATCC 700725) of 15 successive cycles of sub-therapeutic exposures. Bars: Standard error.

**Fig. 7.**
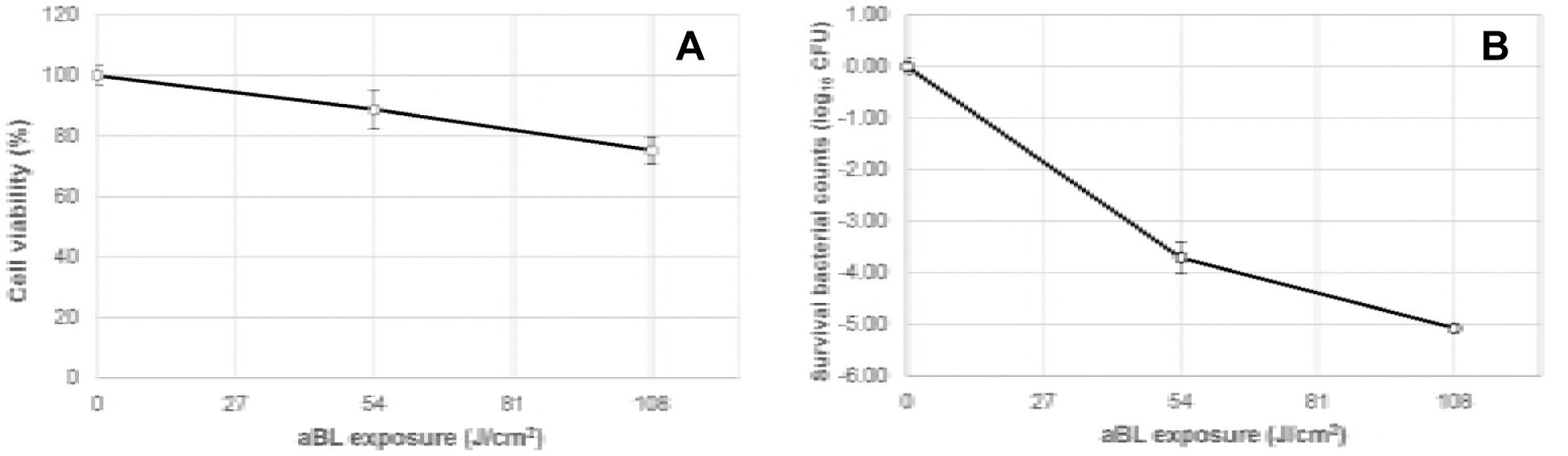
Viability changes of infected human vaginal epithelial cells (VK2/E6E7) by *N. gonorrhoeae* and intracellular *N. gonorrhoeae* in response to 405-nm aBL. **(A)** Infected VK2/E6E7 cells**; (B)** Intracellular *N. gonorrhoeae*. Bars: Standard errors.

The viability loss in the infected VK2/E6E7 cells was 11.3% (0.05-log_10_ CFU) for 54 J/cm^2^ (*P* =0.146) and 24.9% (0.12-log_10_ CFU) for 108 J/cm^2^ (*P* =0.0011), respectively (Fig. 7A). Under the same aBL exposures, 4.18- and 5.07-log_10_ CFU reduction of *N. gonorrhoeae* cells were observed, respectively (Fig. 7B), indicating that *N. gonorrhoeae* cells were still much more susceptible to aBL inactivation than VK2/E6E7 cells in the co-cultures.

Confocal microscope imaging ascertained the invasion of VK2/E6E7 cells by *N. gonorrhoeae* (Fig. 8A). Viable VK2/E6E7 cells and *N. gonorrhoeae* were stained green by SYTO9, and nonviable VK2/E6E7 cells and *N. gonorrhoeae* were stained red by propidium iodide (PI). A few spontaneously dead VK2/E6E7 cells and *N. gonorrhoeae* cells were observed in the culture without aBL irradiation (Fig. 8A).

**Fig. 8.**
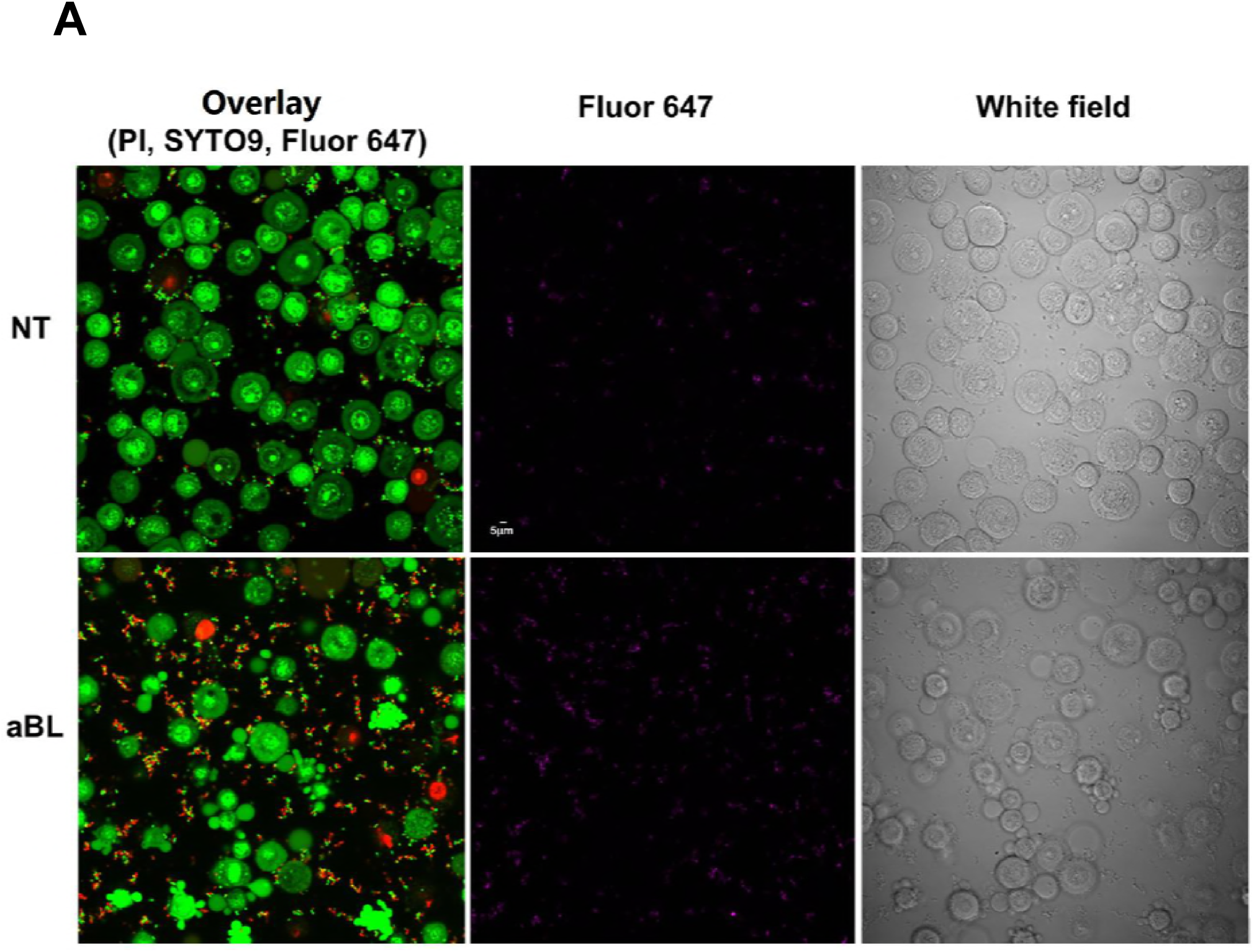
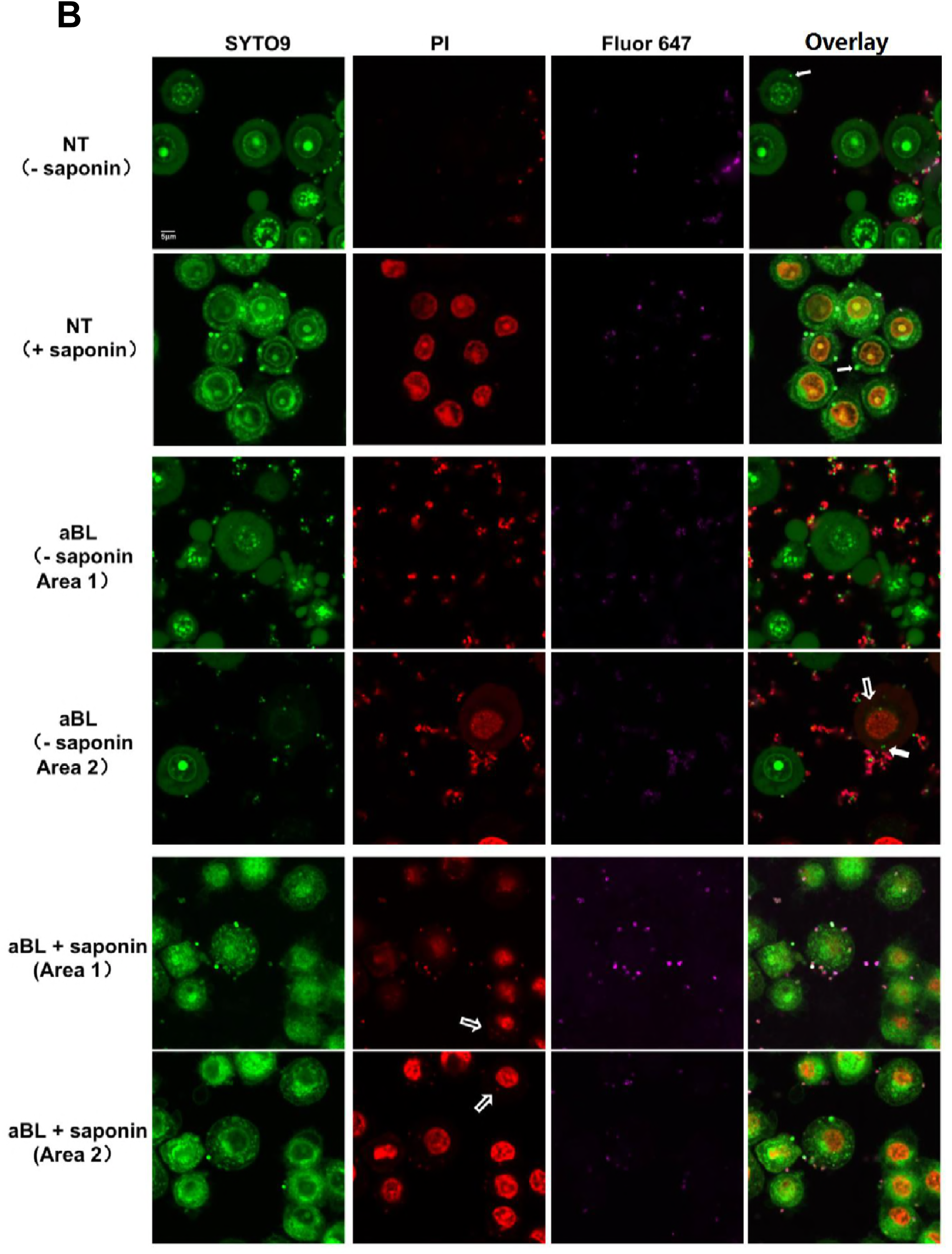
Confocal images of the vaginal epithelial cells (VK2/E6E7) infected with *N. gonorrhoeae* (ATCC700825). (**A**) Low magnification. aBL: treated with 405-nm aBL. 60mW/cm^2^, 108J/cm^2^. NT: no treatment. PI:excitation wavelength 559 nm, emission wavelength 603 nm; SYTO9: excitation wavelength 488nm, emission wavelength 520 nm; Alexa Fluor 647-soybean lectin: excitation wavelength 635nm, emission wavelength 668 nm. Scale bar: 5 µm. (**B**) High magnification. NT: no treatment. aBL: treated with 405-nm aBL (60 mW/cm^2^, 108 J/cm^2^). + saponin: infected cells incubated with 0.05% saponin for 15 min. PI penetrated into the vaginal epithelial cells. - saponin: infected cells not incubated with saponin. External nonviable bacteria exhibit purple + red fluorescence, appear red only; external viable bacteria exhibit purple +green fluorescence; and intracellular viable bacteria exhibit green fluorescence only. Solid white arrows: viable intracellular *N. gonorrhoeae*. Hollow white arrows: Nonviable intracellular *N. gonorrhoeae*. Scale bar: 5 µm.

Confocal microscope imaging also revealed that, after an aBL exposure of 108 J/cm^2^, there was a marked increase in the number of nonviable *N. gonorrhoeae* cells adherent to the VK2/E6E7 cells (Fig. 8A). In contrast, no obvious increase in the number of nonviable VK2/E6E7 cells was observed after aBL (Fig. 8A). For *N. gonorrhoeae* cells, not all of them appeared nonviable immediately after aBL as shown in Fig. 8A. This did not seem consistent with the inactivation results of *N. gonorrhoeae* from colony forming assay, which showed over 5-log_10_ CFU reduction of *N. gonorrhoeae*. A possible reason of this inconsistence is that in addition to cell membrane damage of bacteria, other mechanisms (e.g., oxidative DNA damage) were involved in bacterial cell inactivation without necessarily damaging the cell membrane (36, 37), which were not detectable by PI staining.

Without aBL, under higher magnification (Fig. 8B), viable adherent *N. gonorrhoeae* cells are visible, depicted as green ˝dots˝ by SYTO9. Because *N. gonorrhoeae* are only about 1/10 the size of the vaginal epithelial cells, intracellular viable bacteria could only be identified in certain cross sections of the vaginal epithelial cells (solid white arrows).

In untreated VK2/E6E7 cells, nonviable intracellular *N. gonorrhoeae* cells were rarely observed after permeabilizing the membranes of VK2/E6E7 cells by saponin, as shown by the confocal image (Fig. 8B). When VK2/E6E7 cells (without being permeabilized) were treated with aBL, increased numbers of nonviable adherent *N. gonorrhoeae* cells were observed after aBL exposure. After permeabilizing, nonviable *N. gonorrhoeae* cells (hollow white arrows) were also observed in VK2/E6E7 cells.

## DISCUSSION

In this study, we explored the potential use of aBL for the treatment of gonococcal infection, a disease that is on the rise worldwide, poses an urgent threat to human health, and is becoming untreatable through standard antibiotic strategies. Our findings demonstrated that *N*. *gonorrhoeae*, including antibiotic-resistant strains, were highly susceptible to 405-nm aBL. TEM imaging data suggested that aBL induced damage to the bacterial cell wall and subsequently led to bacterial death. Many studies reported that, in contrast to most other bacterial species, *N. gonorrhoeae* contains no superoxide dismutase (SOD), which is the major antioxidant defense system (38). This might explain the high susceptibility of *N. gonorrhoeae* to aBL inactivation, which is dominantly mediated by singlet oxygen. Unpublished data in our laboratory showed that *N*. *gonorrhoeae* is on average 7.32-, 5.87-, and 28.0-fold more susceptible to aBL than N*Pseudomonas aeruginosa*, *Acinetobacter baumannii*, and *Escherichia coli,* respectively.

We also tested the effectiveness of aBL at another wavelength, 470-nm, for the inactivation of *N*. *gonorrhea*. In contrast to wavelength 405 nm, which targets porphyrins, wavelength 470 nm targets flavins. We found that the anti-gonococcal efficacy of 470-nm aBL was significantly lower than 405-nm aBL. A possible reason for this is the lower singlet oxygen quantum yield of flavins compared with porphyrins upon aBL irradiation (39).

The UPLC analysis in this report confirmed the presence of both endogenous porphyrins and flavins in *N. gonorrhoeae*. The UPLC results also showed that the most abundant type of porphyrin in *N. gonorrhoeae* was coproporphyrin, which is a highly efficient singlet oxygen generator. The singlet oxygen quantum yields was determined to be 0.37 for coproporphyrin III (40). Interestingly, PpIX, which is commonly present in pathogenic bacteria, was not detected in any of the five *N. gonorrhoeae* isolates studied in this report.

The biosynthesis of photosensitizing porphyrins in bacteria occurs in the pathway of the synthesis of heme (41). It is generally believed that the heme synthesis pathway of most bacteria begins with charged glutamyl-tRNA^Glu^ to form the universal precursor ALA, and porphyrins are formed through a series of conserved enzymatic steps. PpIX was believed to be the classical pathway. However, recent studies reported that there is a divergent noncanonical pathway that produces heme through the coproporphyrin III intermediate (42, 43). The lack of endogenous PpIX and rich of endogenous coproporphyrin in *N. gonorrhoeae* indicates that, unlike other Gram-negative bacteria, *N. gonorrhoeae* synthesizes heme via the coproporphyrin III pathway.

In addition to synthesizing its own heme, *N. gonorrhoeae* is able to internalize and utilize exogenous heme within host epithelial cells for growth (44).

The comparison of the aBL-susceptibility between *N. gonorrhoeae* and human vaginal epithelial cells suggested that there is a therapeutic window where *N. gonorrhoeae* could be selectively inactivated while normal vaginal epithelial cells are preserved. This can be explained by our UPLC findings that the content of aBL-active endogenous photosensitizers in the vaginal epithelial cells is much lower than that in *N. gonorrhoeae*.

*N*. *gonorrhoeae* has evolved mechanisms to infect a variety of host cells and subvert clearance by the host immune response. Infection of genital mucosa by *N*. *gonorrhoeae* involves adherence to and invasion of epithelial cells. Therefore, to be an effective therapeutic, aBL must also be able to inactivate intracellular *N. gonorrhoeae*. Our results in this study demonstrated that aBL inactivated *N. gonorrhoeae* both adherent to and inside the vaginal epithelial cells. Although the infection of the vaginal epithelial cells by *N. gonorrhoeae* slightly reduced the tolerance of the vaginal epithelial cells to aBL, we showed that there still exited a therapeutic window of aBL for preferentially inactivate *N. gonorrhoeae* over the vaginal epithelial cells.

The hypothesis that bacteria are unlikely to develop resistance to aBL is supported by the findings in the present study. It is well accepted that photo-oxidative stress reacts with several cellular macromolecules including proteins, lipids, DNA, and RNA, and subsequently results in cell damage (45). This multi-target feature of aBL minimizes the potential of aBL-resistance development by bacteria (21, 22). We experimentally investigated the potential development of aBL-resistance by *N. gonorrhea* by carrying out 15 successive cycles of subtherapeutic aBL inactivation. No development of aBL resistance by *N. gonorrhea* was observed. Our finding in this regard is also consistent with that of a recent study, showing that 15 repeated sub-lethal exposures of *Staphylococcus aureus* suspension did not affect the susceptibility of the organism to 405-nm light (25).

In conclusion, this study provides novel fundamental information regarding the use of aBL for treating gonococcal infection. aBL at 405-nm inactivates antibiotic-resistant *N. gonorrhoeae* while preserving the host cells. Endogenous blue light active photosensitizers (porphyrins and flavins) are present in *N. gonorrhoeae* cells. The development of gonococcal resistance to aBL is unlikely. Taken together, 405-nm aBL is potentially a promising strategy for combating otherwise untreatable multidrug-resistant gonococcal infection. Further studies are warranted to investigate whether other vaginal flora or biofilms might impact the utility of aBL, optimize the aBL parameters, assess the effectiveness and safety of aBL therapy for gonococcal infection in animal models, and explore potential clinical applications.

## MATERIALS AND METHODS

### Blue light sources

Two light-emitting diodes with peak emissions at 405 nm (M405L2, Thorlabs, New Jersey, USA) and 470 nm (M470L2), which are the optimal wavelengths targeting porphyrins (46) and flavins (47), respectively, were used for irradiation. The irradiance of 60 mW/cm^2^, which was optimized based on log_10_ CFU inactivation of bacteria per unit radiant exposure of aBL (J/cm^2^) in a preliminary study, was employed in the study. Light irradiance was measured using a PM100D power/energy meter (Thorlabs, New Jersey, USA).

### *N. gonorrhoeae* strains and culture conditions

ATCC700825 (FA 1090) and four clinical antibiotic-resistant *N. gonorrhoeae* isolates were studied. ATCC700825 was purchased from the American Type Culture Collection (ATCC, Rockville, MD, USA), and the four additional isolates were obtained through the CDC & FDA Antibiotic Resistance Isolate Bank. Bacteria were routinely grown on gonococcal medium base GC (Remel, Lenexa, KS, USA) agar plates containing GCHI enrichment (Remel, Lenexa, KS, USA) and hemoglobin at 37 °C and 5% CO2.

### Human vaginal epithelial cells and growth conditions

Human vaginal epithelial cells, VK2/E6E7 (ATCC^®^ CRL-2616), were purchased from the ATCC (Rockville, MD, USA). Cells were cultured in keratinocyte serum-free medium (Gibco, USA) supplemented with 5 ng/mL recombinant epidermal growth factor, 50 μg/mL bovine pituitary extract (Invitrogen Corporation, Grand Island, NY, USA), and 100 units/mL each of penicillin and streptomycin (Life Technologies, Grand Island, NY, USA) at 37°C and 5% CO_2_. All experiments were performed in the exponential growth phase of VK2/E6E7 cells (48 h after plating).

### aBL inactivation of *N. gonorrhoeae* in suspensions

Overnight *N. gonorrhoeae* cultures were collected from GC agar plates, washed using PBS, and then re-suspended in PBS to OD_600nm_=0.3 (∽10^8^ CFU/mL). Three (3) mL bacterial suspension was added into a 35-mm petri dish prior to aBL exposure. After varying aBL exposures (9, 18, 27, 36, 45, 54, 72, 90, and 108 J/cm^2^) had been delivered, 30-µL samples of the bacterial suspension were taken and *N. gonorrhoeae* CFUs were measured using colony-forming assay. The experiment was performed in 3 independent replicates for each condition. In addition, bacterial suspensions without being exposed to aBL served as the negative controls.

### Evaluation of aBL toxicity to human vaginal epithelial cells

To test the toxicity of aBL to normal vaginal epithelial cells, VK2/E6E7 cells were exposed to aBL at varying exposures of 54 J/cm^2^, 108 J/cm^2^, and 162 J/cm^2^. The viability of VK2/E6E7 cells was measured using MTT assay 24h after aBL exposure. The experiment was performed in 6 independent replicates for each condition.

In addition, aBL-induced DNA damage in VK2/E6E7 cells was evaluated using a CometAssay kit (Trevigen Inc., USA). The experiments were performed based on the Trevigen standard protocol for single cell gel electrophoresis. Briefly, aBL-treated and untreated VK2/E6E7 cells were detached using trypsin. The cell pellets were resuspended at 37 °C with molten LMAgarose after centrifugation and then loaded onto a pre-coated CometSlide^™^ slide (50µL/well), which was immediately cooled on ice. The embedded cells were lysed in cold Trevigen Lysis Solution overnight at 4 °C and then electrophoresed in cold alkaline buffer (pH 13.3). DNA strands/fragments stained with SYBR Gold were visualized under an Olympus FV-1000 confocal microscope at 496/522 nm.

### Ultra-Performance Liquid Chromatography (UPLC) analysis of endogenous aBL-absorptive photosensitizers in *N. gonorrhoeae* and human vaginal epithelial cells

UPLC was used to separate and quantify the endogenous aBL-absorptive photosensitizers (porphyrins and flavins) in *N. gonorrhoeae* and VK2/E6E7 cells. To extract the endogenous photosensitizers from the cells, overnight cultures were washed using PBS. The cell pellets were collected after centrifugation (at 13,500 × g for 6 min), re-suspended in 1.0 mL of extraction solvent (ratio of ethanol to dimethyl sulfoxide to acetic acid, 80:20:1 [vol/vol/vol]), and then stored at -80°C for 24 h. The cell walls were then disrupted by sonication for 20 min. After centrifugation (at 13,500 × g for 6 min), the supernatant was collected as the whole-cell lysates. The protein content in each sample was measured using BCA protein assay.

Standard porphyrins (Chromatographic Marker Kit, Product # CMK-1A, Frontier Scientific, Logan, Utah; and PpIX, Sigma Aldrich) and standard flavins (Sigma Aldrich) were used as the reference compounds. The Chromatographic Marker kit is composed of uroporphyrin I, 7-carboxylporphyrin I, 6-carboxylporphyrin I, 5-carboxylporphyrin I, coproporphyrin, mesoporphyrin IX, mesoporphyrin IX dihydrochloride; and the standard flavins included Flavin mononucleotide(FMN), flavin adenine dinucleotide (FAD), and riboflavin.

UPLC separation and quantitation of porphyrins and flavins in *N. gonorrhoeae* and VK2/E6E7 cells were carried out using a Waters^®^ Acquity UPLC™ System. The system is composed of a binary solvent manager, sample manager, fluorescence detector, column heater and an Acquity UPLC BEH C18, 1.7 µM, 2.1 x 100 mm column. For detecting porphyrins, the excitation was set at 404 nm wavelength and emission at 618 nm; and for detecting flavins, the excitation and emission were set at 260 nm and 470 nm, respectively.

### TEM examination of morphological changes of *N. gonorrhoeae* cells induced by aBL

To examine the morphological changes of *N. gonorrhoeae* cells induced by aBL, untreated and aBL treated ATCC700825 *N. gonorrhoeae* cells were fixed in 1% paraformaldehyde and 1.25% glutaraldehyde immediately after aBL exposures [9 J/cm^2^ (2.5 min illumination), 18 J/cm^2^ (5 min illumination), and 27 J /cm^2^ (7.5 min illumination)] and stored at 4°C for 2 h. The *N. gonorrhoeae* cells were then washed three times with 0.1 M sodium cacodylate buffer after spinning down (1200 rpm, 10 min) and decanting the fixative. Cell pellets were subsequently processed for TEM imaging.

### Assessment of potential development of gonococcal resistance to aBL

To assess if *N. gonorrhoeae* has the potential to develop resistance to aBL, suspensions of ATCC 700825 in PBS were subject to 15 repeated cycles of sub-therapeutic aBL exposures. In the 1st cycle, the aBL exposure was adjusted to produce ∽ 4.0-log_10_ CFU inactivation of *N. gonorrhoeae*, and the same exposure level of aBL was then used throughout the successive cycles. In each cycle, the surviving bacterial cells after aBL exposure were collected, sub-cultured, and grown overnight for the next cycle of aBL inactivation.

### Infection of human vaginal epithelial cells by *N. gonorrhoeae*

*N. gonorrhoeae* have evolved the mechanism to invade into human vaginal epithelial cells to overcome host defense barrier (48). To assess the effectiveness of 405-nm aBL inactivation of intracellular *N. gonorrhoeae*, VK2/E6E7 cells were seeded into a 35-mm petri dish (Transwell^®^, Costar, NY, USA) at a cell density of 2 × 10^5^ cells/dish. The VK2/E6E7 cells were incubated in 2 mL keratinocyte serum-free medium for 48 h at 37 °C. The supernatant was then discarded and 1 mL suspension of ATCC 700825 was added. A multiplicity of infection (M.O.I.) of 50 (bacteria: VK2/E6E7 cells) was used according to the optimized M.O.I. results of a previous study (49). The co-cultures of VK2/E6E7 cells and *N. gonorrhoeae* were incubated at 37 °C with 5% CO_2_ for 4 h. The invasion of *N. gonorrhoeae* cells into VK2/E6E7 cells was examined by confocal microscope (Olympus, FV 1000-MPE Confocal) using the method described previously (50).

### aBL inactivation of intracellular *N. gonorrhoeae*

Four identical cultures (A, B, C, and D) of VK2/E6E7 cells infected with *N. gonorrhoeae* were prepared for each experiment. Culture A was treated with aBL followed by incubation with 0.05% saponin for 20 min to lyse the VK2/E6E7 cells and liberate the bacteria. The viability of *N. gonorrhoeae* was then determined using colony forming assay. Culture B was treated with the same aBL exposure, and the viability of VK2/E6E7 cells was assessed using MTT assay 24 h after aBL exposure. Cultures C and D were not exposed to aBL. The viability of *N. gonorrhoeae* in Culture C was measured using colony forming assay and the viability of VK2/E6E7 cells in Culture D was measured using MTT assay. aBL exposures of 54 J/cm^2^ and 108 J/cm^2^ were tested. The experiment was performed in 6 independent replicates for each condition.

### Fluorescence microscopy for determining the viability of intracellular *N. gonorrhoeae*

For fluorescence microscopy imaging, fluorescent dyes PI and SYTO9 (Invitrogen™) were used to assess the viability of *N. gonorrhoeae* as well as the vaginal epithelial cells immediately after aBL exposure. To discriminate the viability of intracellular bacteria and the bacteria adherent to the vaginal epithelial cells, *N. gonorrhoeae*-infected human vaginal epithelial cells were incubated with Alexa Fluor 647-coupled soybean lectin (Invitrogen™, false-colored purple) for 10 min before adding PI and SYTO9. In the VK2/E6E7 cells with intact cell membranes, intracellular nonviable *N. gonorrhoeae* cells could not be stained by PI because the cell membranes of VK2/E6E7 cells are impermeable to PI. To stain intracellular nonviable *N. gonorrhoeae* using PI, we used 0.05% saponin to permeabilize VK2/E6E7 cells. when adding PI and SYTO9. Under confocal microscopy, nonviable bacteria adherent to the vaginal epithelial cells exhibited purple and red fluorescence, intracellular nonviable bacteria red only, adherent viable bacteria purple and green, and intracellular viable bacteria green only.

### Statistical Analysis

Data were presented as the mean ± standard error. The differences between different conditions were analyzed using one-way analysis of variance (ANOVA). *P* values of <0.05 were considered statistically significant.

## Acknowledgements

### Funding

This study was supported in part by the National Institute Health (R01AI123312 to T.D.) and the U.S. Department of Defense (FA9550-17-1-0277 to T.D.). Y.W. was supported by National Natural Science Foundation of China (61575222) and China Scholarship Council (201603170004).

### Author contributions

Y.W., R.F.E., Y.B., K.D.H., Y.H.G., Y.G., J.A.F., and T.D. designed the experiments. Y.W., R.F.E., Y.B., and X.S.G. performed the experiments. Y.W., Y.B., and T.D. analyzed the data and wrote the manuscript. R.F.E., K.D.H., Y.H.G. and J.A.F. reviewed the manuscript.

### Competing interests

The authors declare that they have no competing interests.

### Data and materials availability

